# BASSL-MI: Batch-Agnostic Self-Supervised Learning Uncovers Clinically Relevant Tumor Niches in Multiplexed Imaging

**DOI:** 10.1101/2025.11.04.686632

**Authors:** Alexander Lin, Shunxing Bao, Ken Lau, Simon Vandekar, Daniel Moyer, Qi Liu, Siyuan Ma

## Abstract

Multiplexed imaging enables rich, high-resolution characterization of the tumor microenvironment but relies on labor-intensive and error-prone cell segmentation and phenotyping pipelines. We present BASSL-MI, a batch-agnostic, self-supervised framework for discovering tissue niches directly from multiplexed imaging data. BASSL-MI operates directly on image patches, eliminating the need for explicit cell segmentation while mitigating image-, sample-, or batch-specific artifacts. Built on a modified contrastive block disentanglement architecture, BASSL-MI learns dual latent representations that separate biologically informative features from batch-dependent factors through spatially guided augmentations and batch-invariance objectives. Applied to a 56-marker colorectal cancer CODEX dataset, BASSL-MI-trained embeddings markedly reduce image-specific variability and recover biologically interpretable spatial niches. Notably, it uncovers CD20–rich follicular regions associated with improved survival, outperforming published findings from cell segmentation-driven clustering. This work demonstrates that self-supervised, patch-based learning can capture clinically relevant spatial organization within tumor microenvironments, advancing toward automated, non-cell-based analysis of multiplexed imaging data.

## 1. INTRODUCTION

Recent advances in multiplexed imaging technologies have transformed our ability to characterize tissue architecture at cellular and subcellular resolution [1, 2, 3]. Unlike traditional immunohistochemistry (IHC), which measures only a handful of protein markers, multiplexed imaging simultaneously quantifies dozens of protein channels [4]. Platforms such as CO-Detection by Indexing (CODEX) [1], Tissue-based Cyclic Immunofluorescence (t-CyCIF) [2], and Imaging Mass Cytometry (IMC) [3] have been successfully applied to diverse health conditions and organs to map protein expression and delineate tissue “niches” – biologically and clinically meaningful organization of cells and protein expression, which have been associated with disease phenotype, risk, and prognosis [1, 5, 6]

Despite these advances, characterizing tissue niches with multiplexed imaging remains challenging because typical workflows depend on error-prone and laborious steps. Analyses commonly begin with cell segmentation, where individual cells are delineated using nuclear and membrane markers. This is complicated by irregularities in cell size and shape (common in tumor tissues) or signal bleed-through [7]. The next step is cell phenotyping, where manual marker gating [8] or clustering approaches adapted from existing single-cell methods [9] are used to assign each cell to a biologically meaningful class. Both methods are labor intensive, require subjective determination of cell phenotypes, and are sensitive to imaging noise and spatial artifacts [10]. Errors in these foundational steps can propagate through subsequent analyses and distort biological conclusions [1, 7], underscoring the need for more robust, automated workflows.

New developments in self-supervised learning (SSL) offer approaches that learn informative embeddings of unlabeled data via pretext tasks, eliminating the need for expert-level annotations [11, 12]. When applied to multiplexed imaging, these approaches can operate on small, multi-channel “patches” cut from larger images [13, 14]. The resulting patch embeddings naturally represent local microenvironments and can be directly used for downstream analysis, bypassing cell segmentation and phenotyping. A key technical challenge, however, is image-specific effects – technical variability arising from differences in e.g. staining, imaging protocols, and sample handling – which, if not corrected for, will dominate patches’ learned representations and obscure true biological signals [15, 16].

Preliminary applications of SSL to multiplexed imaging have shown early success: Atarsaikhan et al. [13] used hierarchical SSL to learn multiscale tissue representations that captured prognostically relevant niches, while Su et al. [14] proposed AdvDINO, a domain-adversarial framework that mitigated slide-specific biases and identified clinically meaningful clusters in lung cancer multiplexed immunofluorescence (mIF) images. However, these methods either do not explicitly address batch effects or rely on adversarial training, which increases computational complexity. Recent work suggests that disentangled or contrastive objectives can achieve batch invariance without adversarial optimization [17, 18], motivating non-adversarial SSL for robust and biologically meaningful representation learning in multiplexed imaging.

We propose BASSL-MI (<u>B</u>atch-<u>A</u>gnostic <u>S</u>elf-<u>S</u>upervised <u>L</u>earning for <u>M</u>ultiplexed Imaging), a framework for clustering tissue niches in multiplexed imaging data. Briefly, our contributions include:

1. Streamlined tissue niche discovery with SSL, operating directly on image patches without cell segmentation/phenotyping steps.
2. Explicit correction of batch effects in multiplexed imaging without adversarial learning, unlike existing methods [13, 14].
3. As validated on a colorectal cancer (CRC) study [19], our learned tumor niches are agnostic to batch effects and discern patient survival, outperforming published, cell-based findings.

## 2. METHODS

### 2.1. Dataset and processing

The dataset used in this work consists of 140 tissue samples collected from 35 CRC patients, each annotated with clinical and demographic metadata such as gender, age, and overall survival. To gain insight into the tumor microenvironment, each tissue sample was imaged with a 56-marker CODEX, encompassing immune, checkpoint, membrane, and other features [19]. Seventeen of the patients exhibited Crohn’s-like reaction (CLR), characterized by the formation of tertiary lymphoid structures (TLSs) – organized immune aggregates that have been associated with improved prognosis [20]. In contrast, the remaining eighteen patients showed diffuse inflammatory infiltration (DII), where immune cells are dispersed throughout the tissue without forming organized structures. This difference in immune spatial architecture has important biological and clinical implications: patients with CLR and TLS formation typically demonstrate significantly better overall survival compared to those with DII, who lack such organized immune responses [19, 20].

Twenty-three of the imaged markers (including HOECHST for nuclei) were selected and utilized in this study, focusing on those that defined cell types and other structural features. To mitigate broader image-specific batch effects, pixel intensity values for these markers were log normalized on a per-channel, per-image basis, following [16]. 128 *×* 128 patches were cut from each image using a sliding window of step size 32 and fed to the self-supervised models for training; the patch dimensions approximately agree with tumor niches as considered in the original publication [19].

### 2.2. Learning framework

BASSL-MI takes inspiration from the supervised contrastive block disentanglement (SCBD) algorithm [18] for SSL. The goal of SCBD is to learn representations of an input *x* that are informative of labels *c* while invariant to nuisance factors *b*. We learn two disjoint embeddings: one encodes information about an entity of interest (*z*_*c*_); the other encodes information about the unwanted factor (*z*_*b*_).

We modify SCBD by generalizing the supervision on *c*. In our formulation, *c* no longer corresponds to a specific class label but instead corresponds to different augmentations (see below) of the same or similar underlying input *x*. This preserves the unsupervised nature of tissue niche discovery while removing unwanted batch effects.

Our method is diagrammed in **Fig. 1**. Given input patches *x*, we learn two encoder networks, *f*_*c*_ and *f*_*b*_, that transform *x* into intermediate representations *z*_*c*_ and *z*_*b*_, respectively.

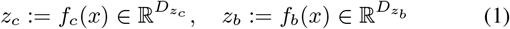

**Fig. 1.**
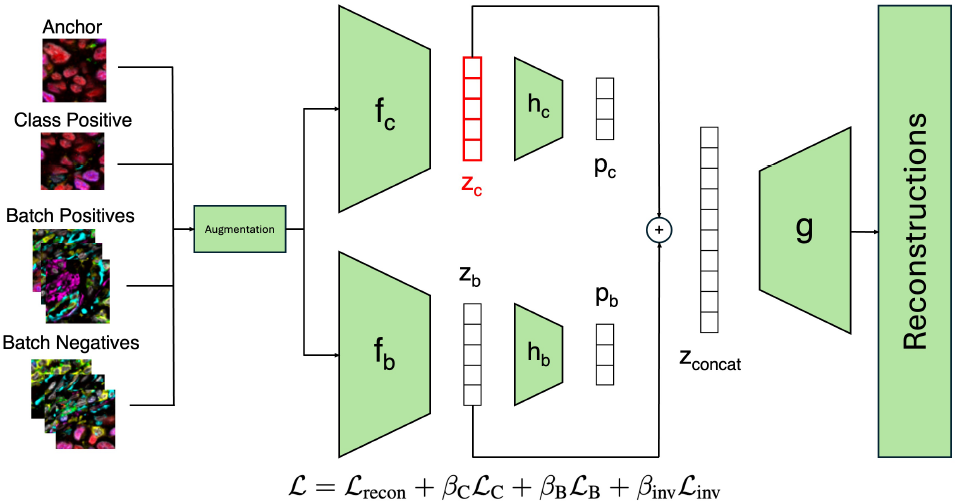
Overview of BASSL-MI. Each anchor patch is a small multi-channel region from a multiplexed image. Augmented neighboring patches serve as class positives (similar biology), same-slide patches as batch positives, and patches from other slides as batch negatives. Two encoders learn separate representations: *f*_*c*_ for biological content (*z*_*c*_) and *f*_*b*_ for batch variation (*z*_*b*_). The decoder reconstructs the input from both latents, while contrastive and invariance losses ensure *z*_*c*_ captures biological instead of batch signals. The red color on *z*_*c*_ indicates this representation is used for downstream analysis.

We also learn two networks, *h*_*c*_ and *h*_*b*_, that map the intermediate *z* representations to their projections, *p*_*c*_, and *p*_*b*_, which are normalized to the unit hyper-sphere and are used for loss function calculations.

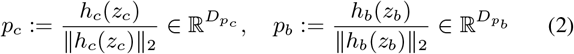

During the decoding process, *z*_*c*_ and *z*_*b*_ are concatenated and passed to the decoder *g* to produce a reconstruction of the input 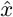.

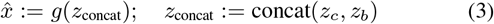

Our objective consists of four components: ℒ_recon_, ℒ _C_, ℒ _B_, ℒ _inv_. Reconstruction loss ℒ _recon_ is the summed mean-squared error between the original input and reconstruction over minibatch ℬ:

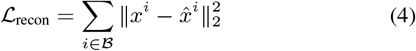

Next, we define the InfoNCE loss [21], which is a major component of the remaining three objectives.

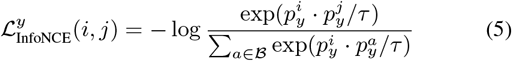

where *y* is the target factor, *τ* is a temperature hyperparameter, *i* is the current sample (anchor) index, and *j* is the positive sample index.

The contrastive loss terms ℒ _C_ and ℒ _B_ are direct applications of the InfoNCE loss as applied to *p*_*b*_ and *p*_*c*_, respectively:

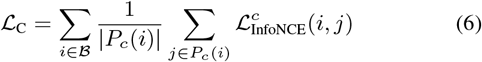

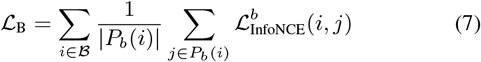

Here, *P*_*c*_(*i*) and *P*_*b*_(*i*) denote the set of positive samples relative to the anchor *i* for factors *c* and *b*, respectively. The final term ℒ _inv_ represents the batch invariance loss. The goal of this objective is to make the *z*_*c*_ representations invariant to the external factor *b*. It performs this optimization by matching the InfoNCE losses on factor *c* between samples within the same batch as anchor *i* and samples from other batches.

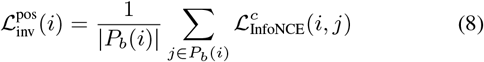

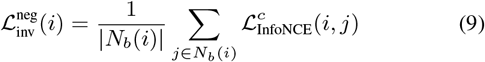

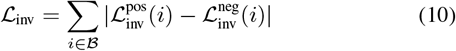

where *P*_*b*_(*i*) is from above and *N*_*b*_(*i*) = ℬ *\*{*P*_*b*_(*i*) ∪ *i* }. From a classification standpoint, this can be seen as minimizing the classification accuracy at the batch level.

The overall objective for this framework can be written as

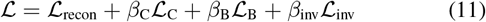

where *β*_C_, *β*_B_, and *β*_inv_ are prespecified hyperparameters.

We implement *f*_*c*_ and *f*_*b*_ as convolutional neural networks followed by fully connected multilayer perceptrons (MLPs) that take flattened convolutional representations and map them to *z*_*c*_ and *z*_*b*_. *h*_*c*_ and *h*_*b*_ are fully connected MLPs, mapping *z*_*c*_ and *z*_*b*_ to *p*_*c*_ and *p*_*b*_. The decoder *g* is a transpose convolutional network that maps the concatenation of *z*_*c*_ and *z*_*b*_ back to the original patch domain.

To complete the contrastive framework, we utilize a spatial heuristic to define positive samples (class positives). For each anchor patch, we select a spatially adjacent patch (or the patch itself) as a positive sample, under the assumption that neighboring regions in tissue share similar biological characteristics. We then apply a series of augmentations – random rotations, horizontal and vertical reflections, and small amounts of Gaussian noise – to both the anchor and positive patches. This spatially informed augmentation strategy encourages the model to learn representations invariant to technical noise and uninformative imaging factors (such as orientation or illumination), while capturing meaningful biological variation.

The external factor *b* that we seek to establish invariance over is the image the patches come from. Previous literature has described the presence of image-specific variations resulting from technical noise, not biological patterns. Thus, we utilize this framework to create batch-agnostic representations *z*_*c*_ and instead store batch-specific information within *z*_*b*_. Patches from the same image serve as batch positives, while patches from other images serve as batch negatives.

### 2.3. Spatially-aware clustering

After model training was complete, we extracted the batch-agnostic latent variables (*z*_*c*_) for all patches in the dataset and performed Leiden clustering [22] on these representations to identify groups of spatially and phenotypically similar tissue regions. Following this, we applied spatial smoothing across neighboring patches prior to analysis to reduce noise from isolated misclustered areas. We then examined marker expression enrichment within each cluster to characterize their underlying biological composition and computed the average cluster frequency per patient to compare the prevalence of specific clusters between the CLR and DII patient subgroups.

### 2.4. Comparison considerations

Of the two existing SSL applications in multiplexed imaging, [13] does not account for image-specific effects and corresponds to a non-batch-adjusted baseline. [14] does not have public code available. As such, we focus our batch correction comparison on non-batch-adjusted baseline and an ablation of our invariance loss component.

We further conduct clinical relevance comparisons of BASSL-MI-identified tumor niches and those based on cell segmentation, phenotyping, and composition clustering (reported in the original publication [19]). Cell-based niche analyses remain the most common analytical approach to date, and comparisons against such cell-based findings provide application baselines for BASSL-MI. Specifically, each patch was assigned a baseline niche label based on annotations from [19], where each cell’s local microenvironment – defined by the composition of surrounding cells – was used to identify its tumor niche. We assigned each patch the plurality tumor niche label among the cells within that patch, breaking ties arbitrarily. These labels were used as a baseline to compare against BASSL-MI-derived niches, in terms of their clinical correlation with patient survival outcome.

## 3. RESULTS

### 3.1. Reduction of image-specific batch effects

BASSL-MI substantially reduces image-specific batch effects in the learned latent representations. As expected, strong batch effects were initially observed across patient images. To overcome this, we incorporated several batch-correction measures into the SSL framework, including blocking the latent space and an explicit batch-invariance loss objective (see Methods for details). Conceptually, the goal of this learning framework is for the model to disentangle biologically relevant features from unwanted variation.

For comparison, we trained a baseline model without the batch invariance loss or partition of the latent space, similar in spirit to [13]. As visualized in **Fig. 2(a)**, the resulting embeddings showed clear batch-specific segregation, with patches clustering primarily by batch in the UMAP. Introduction of the latent space partition disperses the learned representations across the UMAP embedding space, as visualized in panel (b); however, batch-specific artifacts still appear. When latent space blocking is combined with the batch invariance loss, the embeddings become further interleaved across batches (**Fig. 2(c)**), demonstrating increased suppression of batch-driven structure. This trend is consistent with the statistics reported in **Table 1**, where the dependence of the latent space on batch (*R*^2^ ↓) decreases from 0.327 (no correction) to 0.285 (latent space blocking) and further to 0.160 (with invariance loss and blocking), confirming that the combined strategy most effectively mitigates batch effects.

**Table 1.** Ablation Test: Dependency of latent space on image, measured as *R*^2^ from regression of latent dimensions on image.

| Batch Correction Strategy | $R^2 \downarrow$ |
| --- | --- |
| None (baseline) | 0.327 |
| Latent space blocking | 0.285 |
| Latent space blocking + invariance loss | 0.160 |

**Fig. 2.**
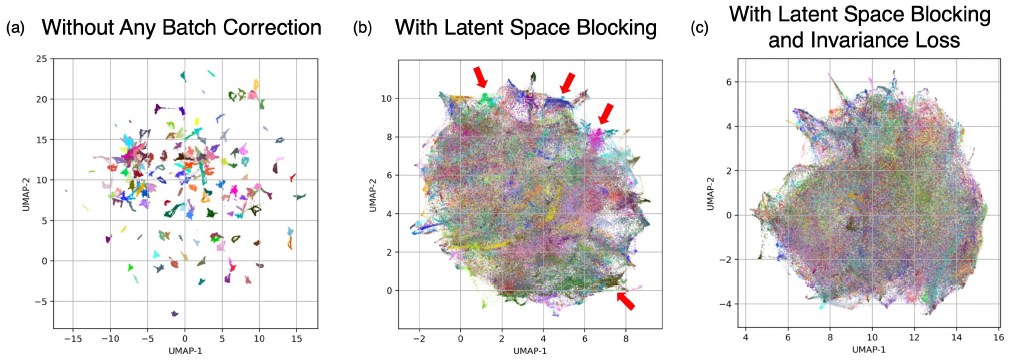
BASSL-MI reduces batch effects in learned representations. Each point represents a patch, colored by its source image (batch). **(a)** Without correction, embeddings cluster strongly by image, indicating pronounced batch effects. **(b)** Latent space blocking reduces this effect, though some batch grouping remains (red arrows). **(c)** Combining latent space blocking with the invariance loss produces well-mixed embeddings, demonstrating that BASSL-MI performs effective batch correction.

### 3.2. Learned latent representations are biologically and clinically informative

BASSL-MI learned tumor niches more truthfully represent protein channels’ spatial organization compared to the published baseline learned through segmented and phenotyped cells (**Fig. 3**). Panel (a) shows two representative images. The left column shows raw marker expression on the tissue; the middle column shows the distribution of BASSL-MI niches; and the right column shows the distribution of the baseline niche labels derived from [19] (cell-based clusters). Visually, for the top sample, BASSL-MI clusters define the major follicular region in the center of the image. The four epithelial regions on the left, which overlap with the MUC-1 marker in the original image, are identified in the four red globular regions from BASSL-MI latent representations (cluster 5). The yellow niche (cluster 7) corresponds with the large smooth muscle region at the top of the image, and the magenta niches (cluster 10) correspond with increased expression of CD56, a marker for NK cells. In contrast, the tumor niches from [19] capture some of these regions, but they notably miss the NK cell-enriched regions and some of the epithelial regions. For the second sample, we see increased CD20 expression (in green) in the top of the original image. This sample is from a CLR patient, indicating a likely representation of a TLS. BASSL-MI representations successfully recaptures this (cluster 20), whereas the baseline labels classify that region as T-cell enriched.

**Fig. 3.**
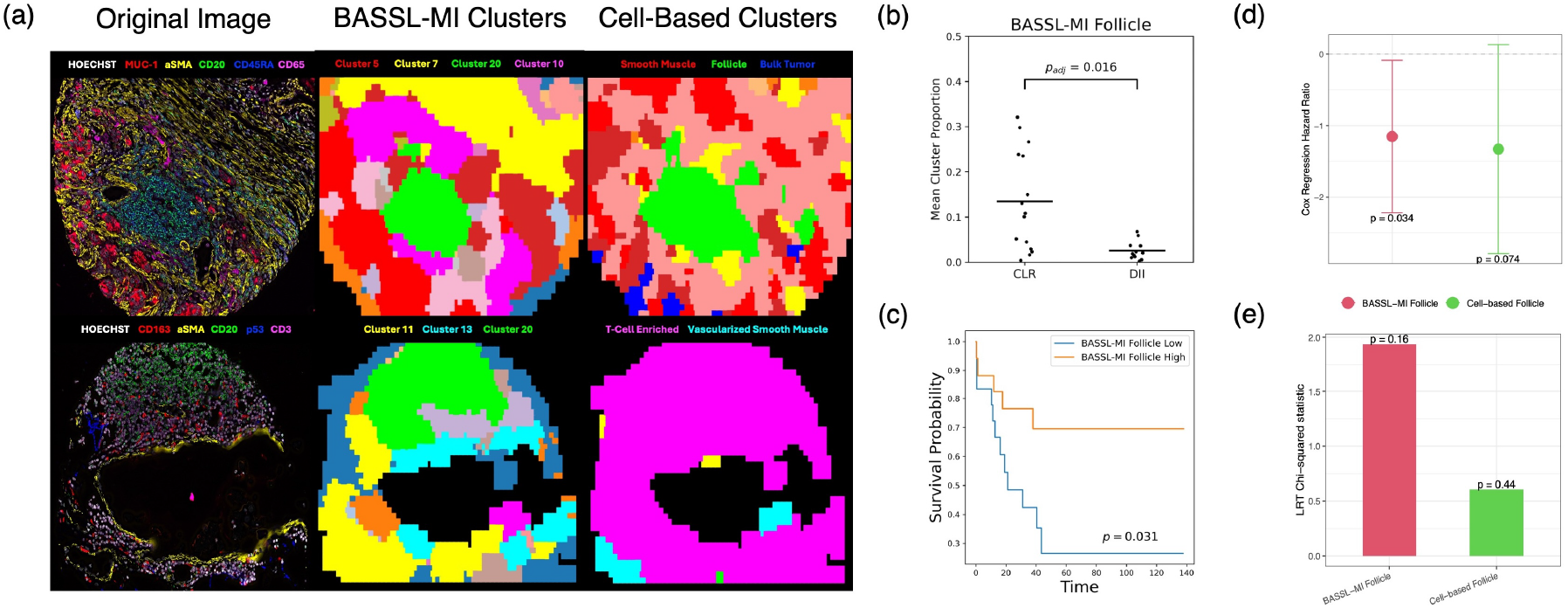
Biological and clinical relevance of BASSL-MI tumor niches. **(a)** Representative visualization of learned niche labels demonstrate better agreement between BASSL-MI-identified niches and actual protein expression, compared to published, cell-based labels [19]. **(b)** The BASSL-MI Follicle niche is enriched in CLR compared to DII patients, aligning with expectations. **(c)** Frequency of BASSL-MI Follicle niches is associated with overall survival. Kaplan-Meier survival curves represent patient stratification using the niche frequency’s median. **(d)** When separately associated with patient survival with Cox proportional hazard regression, frequency of BASSL-MI follicle had a significant hazard ratio whereas cell-based follicle did not. **(e)** In joint likelihood ratio (LRT) analysis of survival versus the two follicle labels, BASSL-MI follicle had a higher *χ*^2^ statistic, indicating a stronger signal for survival compared to cell-based follicle when mutually adjusted.

The BASSL-MI-identified B-cell niche has clinically meaningful associations with patient phenotype and outcome (**Fig. 3(b,c)**). Our cluster 20 exhibited notably high expression of CD20, a canonical B-cell marker, suggesting correspondence to B-cell–rich follicular regions (present in CLR patients, largely absent in DII patients). We thus term it as “BASSL-MI follicle” and further examine its validity and clinical relevance. First, in terms of the proportion of the niche in each patient, it was significantly enriched in CLR patients (*p*_adj_ = 0.016, Benjamini–Hochberg correction [23], **Fig. 3(b)**), congruent with expectations. Furthermore, Kaplan–Meier survival analysis based on stratification by median niche frequency (high versus low) demonstrated a significant difference in overall survival (*p* = 0.031, **Fig. 3(c)**). These findings agree with [19] and validate BASSL-MI’s ability to identify distinct spatial features of CLR tissues and their associations with improved clinical outcomes.

Furthermore, BASSL-MI-identified follicles have improved discrimination of patient survival compared to baseline, cell-based follicles (**Fig. 3(d**,**e)**). We fit marginal Cox proportional hazard regressions associating patient survival with the frequency of either BASSL-MI follicles or cell-based follicles. The hazard ratio is significant for BASSL-MI follicle frequency (*p* = 0.034), but non-significant for cell-based follicle (**Fig. 3(d)**). We further conduct likelihood ratio testing (LRT) by comparing the above marginal regressions against a mutually adjusted Cox regression with both terms included. While neither is significant, removing BASSL-MI follicle frequencies from the full model yields a higher *χ*^2^ statistic (*p* = 0.16), suggesting a trend that BASSL-MI follicles are more informative for survival (**Fig. 3(e)**). These findings suggest that follicle structures identified by BASSL-MI have closer associations with patient survival compared to baseline cell-based labeling, likely due to BASSL-MI’s more truthful alignment between identified tumor niches and actual protein expression in the original samples.

## 4. CONCLUSION

Existing multiplexed imaging analysis relies on manual and sensitive algorithms for cell segmentation and phenotyping, where errors can propagate and potentially distort conclusions. Moreover, these approaches reduce rich spatial information to the single-cell level, losing critical tissue contexts. We present BASSL-MI, a batch-agnostic, self-supervised framework that does not require cell segmentation/phenotyping, corrects for batch effects inherent to multiplexed imaging, and produces biologically interpretable niches. It represents one of the first steps toward non-cell-based multiplexed imaging analysis. Future extensions will explore more disease types, semi-supervised strategies to incorporate human annotation, and higher-order spatial interactions between tissue niches.

## 5. COMPLIANCE WITH ETHICAL STANDARDS

This work analyzes public, de-identified tissue section imaging data and represents secondary use of existing, non-identifiable research data. No new human subjects were recruited, and no personally identifiable information was accessed or analyzed. As such, it was exempt from institutional review board (IRB) oversight.

## 6. ACKNOWLEDGMENTS

This work is partially supported by the Vanderbilt-Ingram Cancer Center GI SPORE Career Enhancement Program (parent grant NIH U2CCA233291). The authors would like to acknowledge input from Dr. Lianrui Zuo and Dr. Matthew Berger in supporting this work.

## Notes

### Competing Interest Statement

The authors have declared no competing interest.

